# Judgments of risk of bias associated with random sequence generation in trials included in Cochrane systematic reviews are frequently erroneous

**DOI:** 10.1101/366674

**Authors:** Ognjen Barcot, Matija Boric, Tina Poklepovic Pericic, Marija Cavar, Svjetlana Dosenovic, Ivana Vuka, Livia Puljak

## Abstract

**Background:** Purpose of this study was to analyze adequacy of judgments about risk of bias (RoB) for random sequence generation in Cochrane systematic reviews (CSRs) of randomized controlled trials (RCTs).

**Methods:** Information was extracted from RoB tables of CSRs using automated data scraping. We categorized all comments provided as supports for judgments for RoB related to randomization. We analyzed number and type of various supporting comments and assessed adequacy of RoB judgment for randomization in line with recommendations from the Cochrane Handbook.

**Results:** We analyzed 10527 RCTs that were included in 729 CSRs. For 5682 RCTs randomization was not described; for the others it was indicated randomization was done using computer/software/internet (N=2886), random number table (N=888), mechanic method (N=366), or it was incomplete/inappropriate (N=303).

Overall, 1194/10125 trials (12%) had erroneous RoB judgment about randomization. The highest proportion of errors was found for trials with high RoB (28%), followed by those with low (19%), or unclear (3%). Therefore, one in eight judgments for the analyzed domain in CSRs was erroneous, and one in three if the judgment was “high risk”.

**Conclusion:** Cochrane systematic reviews cannot be necessarily trusted when it comes to judgments for risk of bias related to randomized sequence generation.

## Background

Randomized controlled trials (RCTs) are considered crucial for objectively testing efficacy and safety of interventions [1]. They are often used to inform clinical practice and included in systematic reviews to obtain even higher level of evidence through evidence synthesis. However, recent systematic review of meta-epidemiological studies indicated that estimates about effects of interventions might be exaggerated in RCTs with inadequate or unclear sequence generation and blinding [2].

RCTs can, therefore, be biased if flaws in design and conduct lead to overestimation or underestimation of intervention effect estimates. Because of this potential for bias, assessment of the risk of bias (RoB) in trials is one of the usual methodological steps during preparation of Cochrane systematic reviews (CSRs). Bias has been defined as any systematic error that can negatively impact estimated effects of interventions and lead to wrong conclusions about efficacy and safety of analyzed interventions [3].

In Cochrane’s RoB tool systematic review authors are expected to provide judgment about the level of RoB as low, unclear, or high for seven different potential domains of bias. Cochrane authors should also provide accompanying comment that needs to justify the judgment. The first domain analyzed in the Cochrane RoB tool is assessing randomization sequence as a part of selection bias [4].

A recent report analyzed the evolution of reporting and inadequate methods over time in 20920 RCTs included in Cochrane reviews [5]. The authors analyzed data from RCTs included in all CSRs published between March 2011 and September 2014, which reported an evaluation of the Cochrane RoB items, including sequence generation. The results indicate that unclear risk for sequence generation was found in 49% of trials, high risk in 4% of all trials, and low risk in 48% of trials included in analyzed CSRs [5]. Additionally, it was found that the proportion of trials with unclear RoB for sequence generation decreased over time, falling from 69% in 1986-1990 cohort of trials to 31% in 2011-2014 cohort. The proportion of trials with high RoB for sequence generation fell from 4.6% in 1986-1990 to 3.2% in 2011-2014 [5].

However, it is possible that the way the Cochrane authors judge RoB for sequence generation is highly variable. We have already proved this for RoB domains for attrition bias and other bias (Babic et al, unpublished data). In that case, results presented in the study of Dechartres et al. [5] or similar studies, would not be based on consistent ratings of RoB in Cochrane reviews, and improvements shown for certain RoB domains could be misleading.

The objectives of this study were two-fold: to evaluate the rationales based on which judgments related to random sequence generation were made for trials in Cochrane reviews, and to investigate the proportion of erroneous judgments about randomization based on independent re-assessments using guidance from the Cochrane Handbook.

## Methods

### Study design

This was a meta-epidemiological study that analyzed methods of published CSRs.

### Inclusion and exclusion criteria

We retrieved CSRs of RCTs about interventions published from July 2015 to June 2016 (N = 955) from The Cochrane Library via advanced search. We excluded all CSRs (N=226) that did not include RCTs about interventions (diagnostic CSRs, overviews of systematic reviews), as well as empty reviews, and reviews that were withdrawn in the analyzed period. If a CSR included both randomized and non-randomized trials, we analyzed RoB table only for included RCTs.

### Screening for study eligibility

One author assessed all titles/abstracts to establish eligibility of CSRs for inclusion. The second author verified assessments of the first author.

### Data extraction

Data extraction was automated in a stepwise manner in MS Excel 2010 (Microsoft, Redmond, WA, USA) using macro-commands written in VBA (Visual Basic for Applications, Microsoft, Redmond, WA, USA) by author OB. Data scraping was done by automated copying of all the content from The Cochrane Library webpage for every eligible CSR to a separate spreadsheet in MS Excel. Raw data were trimmed, filtered, and RoB tables extracted for every study included in a CSR by a series of macro-commands. Excel spreadsheet was created containing all the details of random sequence generation domain extracted from corresponding RoB table for every included study.

### Calibration of categorizations

First author (OB) analyzed first 1500 trials against categories provided in the Cochrane Handbook, which was verified by the last author (LP). This calibration exercise was used to create a spreadsheet with drop-down menus that included pre-determined categories. Further categorizations were conducted by four persons (MB, TPP, MC, SD), and then verified by OB. All authors involved in categorizations are medical doctors familiar with Cochrane RoB tool.

### Outcomes

We analyzed number and type of various supporting comments for the Cochrane RoB domain of random sequence generation. We also analyzed adequacy of judgments for this domain, in line with recommendations from the Cochrane Handbook. The Handbook was used as a gold standard in our assessment; if the judgments of Cochrane authors were contrary to the guidance from the Cochrane Handbook, we considered them erroneous.

### Categorizations

To categorize supports for judgments, we used recommendations from the Cochrane Handbook’s Table 8.5.d: Criteria for judging the risk of bias in the “Risk of bias” assessment tool. Firstly, we categorized all the supports for judgments into 12 categories depending on the comment that the Cochrane authors provided to explain their judgment (Table 1). For the low risk of judgment we used all seven examples from this tool as independent categories supporting the judgment (random number table, computer random number generator, coin tossing, shuffling cards or envelopes, throwing dice, drawing of lots, and minimization). Additionally, we created a separate category for comments stating a webpage was used as a method of randomization where http address was cited, and one for Interactive Voice Response System (IVRS).

**Table 1.** Modified Cochrane Handbook’s Table 8.5.d

| **Level of bias** | **Judgments used in this study** |
| --- | --- |
| Criteria for a judgment of "Low risk" of bias. | The investigators describe a random component in the sequence generation process such as:<br>• Referring to a random number table<br>• Using a computer random number generator<br>• Coin tossing<br>• Shuffling cards or envelopes<br>• Throwing dice<br>• Drawing of lots<br>• Minimization<br>+ Web based*<br>+ IVRS<br>+ Random number generator (without mention of a computer) |
| Criteria for the judgment of "High risk" of bias. | The investigators describe a non-random component in the sequence generation process. Usually, the description would involve some systematic, non-random approach:<br>• Quasi random<br>Other non-random approaches happen much less frequently than the systematic approaches mentioned above and tend to be obvious. They usually involve judgment, or some method of non-random categorization of participants:<br>• Judgment, preference, availability<br>+ Incomplete, erroneous randomization |
| Criteria for the judgment of "Unclear risk" of bias. | Insufficient information about the sequence generation process to permit judgment of "Low risk" or "High risk".<br>+ Not described<br>+ Block randomization/stratification<br>+ Envelopes<br>+ Central randomization (statistics department, pharmacy, third party)<br>+ Baseline imbalance between groups<br>+ Baseline balance between groups |

\*Categories added to original table bulleted by plus sign. †IVRS = Interactive Voice Response System.

According to the Cochrane Handbook, additional two categories have been made supporting high risk judgment: quasi random for every type of randomization using some form of systematic non-random component, and randomization according to the judgment of a physician, preference of a subject, or availability of intervention. We added another high risk judgment category – incomplete or erroneous randomization when comments indicated that randomization was partial or flawed in any way.

For every comment not permitting categorization to any of the 12 high or low risk comment categories we created a new category – method of randomization was not described. This included all the instances when the Cochrane authors indicated that something was not described, or have only indicated that the study was described as randomized, without mentioning a method of randomization.

“Method of randomization was not described” category was also used when, in supporting judgments, we found descriptions such as, quote: *no information, no information available, not described, not stated, not reported, unreported*, and similar. In such cases, we assumed that the review authors wanted to say that the method of randomization was not described. Similar to this, “method of randomization was not described” category was used if the Cochrane authors wrote only that information about randomization was on certain page/table/figure, but without any details what is written there; comments such as, quote: *assumed*, *same as above, as above, see previous, see XY study, appendix*, *used CONSORT flow diagram*, *Chinese article, translation required*, *trial was stopped*, *not adequately designed, study withdrawn prior to enrollment*.

Further on, the following supporting comments were categorized as “method of randomization was not described” if the support for judgment mentioned only: baseline characteristics of participants; block randomization or stratification; randomization by envelopes; central randomization by statistics department, pharmacy, or third party without a description of a method used. Based on these supporting comments we created five subcategories in the “method of randomization was not described” parent category. Some of these supporting comments indicate certain aspects of methodology associated with random sequence generation, but not sufficiently enough to be properly judged.

If the Cochrane authors only wrote in a support for judgment that “random number generator” was used, without mentioning computer, this was put in a separate category – “random number generator (without mention of computer)”.

Thus, we created a total of 19 categories of supports for judgment, which were grouped into the following 5 parent categories: i) method of randomization was not described, ii) random number table, iii) incomplete or inappropriate randomization, iv) randomization via computer/software/internet that precludes use of electronic automation as in IVRS or random number generation or use of complex algorithms as in minimization, v) mechanic method of randomization such as coin tossing, drawing of lots, shuffling cards or envelopes and throwing dice.

### Statistics

We used frequencies and percentages to present descriptive data. Subgroup analysis was conducted for CSRs with different types of therapies.

## Results

We analyzed 10527 RCTs included in 729 CSRs. We excluded 402 RCTs from the main analysis because there was no domain for random sequence generation in the RoB table, the supporting comment only indicated “not applicable” or “N/A”, there was no support for judgment, or the study was not an RCT (Table 2). All those studies were checked to ensure that they indeed belong to studies that had randomized design, but we excluded them because we had no way of knowing what the Cochrane authors meant by “not applicable” or “N/A”.

**Table 2.** Reasons for exclusion of trials from analysis

| **Reason for exclusion** | **Total, N (%)** | **High, N (%)** | **Unclear, N (%)** | **Low, N (%)** |
| --- | --- | --- | --- | --- |
| There was no domain for bias associated with random sequence generation in the Risk of Bias table | 218 (54) | Not applicable – no domain |  |  |
| "N/A"* | 121 (30) | 79 (65) | 42 (35) | 0 (0) |
| "not an RCT" † | 49 (12) | 24 (49) | 13 (26) | 12 (25) |
| There was no support for judgment for this domain, only judgment | 14 (4) | 0 (0) | 1 (7) | 13 (93) |
| Total | 402 (100) | 103 (56) | 56 (30) | 25 (14) |

\*N/A = not applicable. †RCT = randomized controlled trial.

### Number and type of various supporting comments

In our main analysis, we categorized support for judgment in the remaining 10125 RCTs into five categories. By using categories from Cochrane Handbook, in more than half of those RCTs method of randomization was not described (N=5682), while the remaining supporting comments indicated that the randomization was done using computer/software/internet (N=2886), random number table (N=888), mechanic method of randomization (N=366), and incomplete or inappropriate randomization (N=303). The frequency of these categories, and different types of supporting comments, which we found in each of these five categories, are presented in Table 3.

**Table 3.** Supporting comments for judgment of risk of bias regarding randomization

| **Categories of supporting comments for judgments of bias associated with randomization** | **Total, N (%)** | **High, N (%)** | **Unclear, N (%)** | **Low, N (%)** |
| --- | --- | --- | --- | --- |
| **Method of randomization was not described** | **5682 (56)** | **80 (1)** | **4655 (82)** | **947 (17)** |
| not described | 4582 (81) | 67 (2) | 4275 (93) | 240 (5) |
| block randomization/stratification | 684 (12) | 2 (0) | 220 (32) | 462 (68) |
| envelopes | 246 (4) | 3 (1) | 117 (48) | 126 (51) |
| central randomization (statistics department, pharmacy, third party) | 99 (2) | 1 (1) | 0 (0) | 98 (99) |
| baseline imbalance between groups | 42 (1) | 7 (17) | 35 (83) | 0 (0) |
| baseline balance between groups | 29 (0) | 0 (0) | 8 (28) | 21 (72) |
| **Random number table** | **888 (9)** | **11 (1)** | **31 (4)** | **846 (95)** |
| random number table | 888 (100) | 11 (1) | 31 (4) | 846 (95) |
| **Incomplete or inappropriate randomization** | **303 (3)** | **260 (86)** | **32 (10)** | **11 (4)** |
| quasi random | 243 (80) | 211 (86) | 23 (10) | 9 (4) |
| incomplete, erroneous randomization | 34 (11) | 28 (82) | 6 (18) | 0 (0) |
| judgment, preference, availability | 26 (9) | 21 (80) | 3 (12) | 2 (8) |
| **Randomization via computer/software/internet** | **2886 (29)** | **2 (0)** | **36 (1)** | **2848 (99)** |
| computer / software generated | 2474 (85) | 0 (0) | 17 (1) | 2457 (99) |
| IVRS* | 60 (2) | 0 (0) | 7 (12) | 53 (88) |
| minimization | 111 (4) | 2 (2) | 8 (7) | 101 (91) |
| random number generator (without mention of computer) | 111 (4) | 0 (0) | 0 (0) | 111 (100) |
| web based | 130 (5) | 0 (0) | 4 (3) | 126 (97) |
| **Mechanic method of randomization** | **366 (4)** | **7 (2)** | **37 (10)** | **322 (88)** |
| coin tossing | 85 (23) | 0 (0) | 1 (1) | 84 (99) |
| drawing of lots | 209 (57) | 4 (2) | 12 (6) | 193 (92) |
| shuffling cards or envelopes | 59 (16) | 3 (5) | 24 (41) | 32 (54) |
| throwing dice | 13 (4) | 0 (0) | 0 (0) | 13 (100) |
| **Total, N (%)** | **10125** | **360 (4)** | **4791 (47)** | **4974 (49)** |

10125 trials included in the main analysis of eligible Cochrane systematic reviews. \*IVRS = Interactive Voice Response System.

### Adequacy of judgment for risk of bias associated with random sequence generation

The majority of the five parent categories of supporting comments had correct judgments by the Cochrane authors. In the category where the method of randomization was not described, 82% of trials were correctly judged as unclear. For random number table 95% of Cochrane authors correctly judged this as low risk of bias. When the Cochrane authors indicated that there was incomplete or inappropriate randomization, 86% of such trials were correctly judged as having a high risk of bias associated with random sequence generation. For supporting comments describing randomization via computer/software/internet, 99% of the authors correctly judged it as low risk of bias. For mechanic methods of randomization, 88% of trials were correctly judged as having a low risk of bias.

Overall, 1194/10125 trials (12%) were erroneously judged for risk of bias associated with random sequence generation. The highest proportion of mistakes was observed in the parent category “method of randomization not described”, where 99% of the trials where supporting comment only indicated that there was central randomization, but the method of randomization was not described, judged it erroneously as low risk of bias. Likewise, 72% of trials that only wrote about baseline balance between groups, 68% of the trials that had only description of block randomization or stratification, and 51% of trials for which supporting comment only mentioned that randomization was done using envelopes were judged as having a low risk of bias (Table 3).

The highest proportion of errors was found in RCTs categorized as having a high risk of bias associated with random sequence generation. There were 360 (3.5%) of RCTs judged as having a high risk of bias for this domain, of which 100 (28%) were erroneously judged. Of the five parent categories we used, only “category incomplete or inappropriate randomization” should have been judged as high risk of bias.

Among the RCTs judged as having a low risk of bias for random sequence generation (N=4974), there were 958 (19%) with erroneous judgment. Among the five parent categories, low risk of bias should be associated with using random number table, randomization via computer/software/internet, and mechanic methods of randomization.

There were 4791 RCTs judged as having an unclear risk of bias for random sequence generation, of which 136 (3%) erroneously judged. Of the five parent categories we used, only comments in the category where the method of randomization was not described should have been judged as having an unclear risk of bias.

## Discussion

Analysis of judgment and comments about the risk of bias associated with random sequence generation in 10125 RCTs included in 729 CSRs indicated that Cochrane authors do not adhere to recommendations regarding assessment of this particular type of risk of bias, and that 12% of judgments for bias associated with this domain were erroneous. This means that one out of every eight judgments of bias regarding randomization of participants in Cochrane reviews is wrong or not elaborated. The most common errors were observed for category of trials judged as having high RoB for sequence generation, where 28% of the judgments were not supported with the explanations given in the accompanying supporting comments for the judgment. To our best knowledge, this is the first meta-epidemiological study evaluating rationales for and accuracy of judgments for risk of bias associated with random sequence generation in Cochrane systematic reviews.

The most frequent error was judging RoB for randomized sequence generation as low, but without sufficient information about the method of randomization. The most frequent such supporting comments were related to mentions of central randomization without further details, baseline balance between the groups where the Cochrane authors assume that the randomized sequence generation was adequate, comments about block randomization or stratification, and supporting comments about using envelopes without further details.

Cochrane Handbook explicitly warns: *“Sometimes trial authors provide some information, but they incompletely define their approach, and do not confirm some random component in the process. For example, authors may state that blocked randomization was used, but the process of selecting the blocks, such as a random number table or a computer random number generator, was not specified. The adequacy of sequence generation should then be classified as unclear”* [6].

The Cochrane is currently undertaking development of *A revised tool to assess risk of bias in randomized trials* (RoB 2.0) [7]. In the RoB 2.0 version baseline characteristics of participants are featured prominently. The RoB 2.0 uses baseline imbalances to signal problems with the randomization process, and one of the signaling questions in the RoB domain about randomization process is: “*Were there baseline imbalances that suggest a problem with the randomization process?*” [7].

We found 111 instances of random number generation were mentioned without remarks about using a computer, electronic calculators, etc. This category was judged as low risk in 100%, and we left it in computer parent category although it did not fulfill strict Cochrane Handbook rules. Otherwise erroneous levels would rise from 12% to 13% in total and from 19% to 22% for trials judged as having a low risk of bias.

Our findings have a profound effect on the reliability of conclusions in Cochrane reviews, and a number of meta-epidemiological studies that were based on the RoB assessments from Cochrane reviews. RoB assessment is regularly mentioned in conclusions of Cochrane reviews. For example, our data indicate that every third judgment indicating that a trial included in a Cochrane review has a high risk of bias associated with random sequence generation is erroneous.

It is very common in Cochrane reviews to read that included studies were of high risk of bias, and therefore their results are less reliable, and we need new, high quality trials. However, if those conclusions are supported with RoB assessment where one in five judgments is erroneous, then the conclusions of Cochrane reviews are severely compromised. Therefore, having the highest number of errors in the group of CSRs that were judged as having low RoB implies that potentially many recommendations indicating that evidence base is worrying, because the readers will get the message that certain recommendations rely on evidence with low RoB, and therefore evidence that is more reliable and trustworthy.

In previous studies authors appear to automatically assume that RoB judgments in Cochrane reviews were correct, and various conclusions were reached, related to those judgments [5, 8]. A recent study reported that poor reporting and inadequate methods have decreased over time, particularly for sequence generation and allocation concealment, based on the analysis of 20920 RCTs included in CSRs [5]. However, our study indicated that this result does not have to be due to better reporting and better methods, but errors in judgment of Cochrane authors.

In our study we found that more than half of the trials included in our sample of CSRs had unclear risk of bias for generating randomization sequence, which is in line with a report of Kahan et al, who reported that risk of selection bias is difficult to ascertain in the majority of trials because of poor reporting [9].

Likewise, studies that analyzed the association between RoB and effect of interventions could have reached erroneous conclusions because they trusted the judgment of systematic review authors. Savovic et al. reported that estimates of intervention effect were exaggerated by the average of 11% in clinical trials with inadequate or unclear sequence generation. Their results were based on analysis of 1973 trials included in 234 meta-analyses [8].

According to the Cochrane Handbook, Cochrane authors can make assumptions, but need to elaborate them [4]. We found supporting comments where Cochrane authors just wrote *assumed*, and judged the trial as having low risk of bias related to randomization, without explaining why they assume that risk is low.

A limitation of our study is confined time period in which analyzed Cochrane reviews were published. However, we analyzed a high number of Cochrane reviews, with a high number of included trials, and these Cochrane reviews were published recently. Therefore, we believe that they are representative of the current state of reporting of the analyzed domain in Cochrane systematic reviews.

It has already been shown that the Cochrane RoB has low reliability between individual reviewers, as well as across consensus assessments of reviewer pairs [10]. Da Costa et al. have argued that low reliability of the RoB assessment in systematic reviews can have detrimental effects on decision making and healthcare quality [11]. Interventions such as standardized intensive training on RoB assessment were tested, and the results indicate that such interventions can improve significantly reliability of the Cochrane RoB tool [12]. Apart from author training, other solutions for improving reliability of RoB judgments would be more stringent peer review and editorial assessment of judgments and supporting comments in the Cochrane RoB table.

Additionally, we analyzed only Cochrane reviews, which use Cochrane RoB tool. Recent studies have indicated that the RoB instrument did not adequately capture risk of bias in RCTs [13, 14]. Future studies on Cochrane RoB 2.0 tool are warranted to see how the new tool compares to the current one.

## Conclusion

Cochrane systematic reviews cannot be necessarily trusted when it comes to judgments for risk of bias related to randomized sequence generation, particularly when the Cochrane authors judge that the risk is high. Interventions are necessary to improve reliability of Cochrane risk of bias assessment.

**List of abbreviations**
CDSR: – The Cochrane Database of Systematic Reviews
CONSORT: – Consolidated Standards of Reporting Trials
CSR: – Cochrane Systematic Review
et al.: – *et alia:* and others
IVRS: – Interactive Voice Response System
RCT: – Randomized Controlled Trial
RoB: –Risk of Bias
VBA: – Visual Basic for Applications

## Declarations

### Ethics approval and consent to participate

This study involved only analysis of data from published scientific literature; we did not collect any primary data.

### Consent for publication

Not applicable.

### Availability of data and materials

Analyzed data were publicly available information availabile in the risk of bias tables of Cochrane systematic reviews published in the Cochrane Database of Systematic Reviews (CDSR), http://www.cochranelibrary.com. The datasets used and/or analysed during the current study are available from the corresponding author on reasonable request.

### Authors’ contributions

Study design: LP, data extraction: OB, IV, data analysis and interpretation: OB, MB, TPP, MC, SD, IV, LP, writing the first draft of the manuscript: OB, LP. Revisions of the manuscript for important intellectual content: OB, MB, TPP, MC, SD, IV, LP, final approval of the manuscript: OB, MB, TPP, MC, SD, IV, LP, agree to be accountable for all aspects of the work: OB, MB, TPP, MC, SD, IV, LP. Guarantor: LP.

## Acknowledgements

Not applicable.

